# Prediction in trait-based ecology: global simulations of specific leaf area using a trait-based dynamic vegetation model

**DOI:** 10.1101/2025.04.14.648688

**Authors:** Liam Langan, Teja Kattenborn, Christine Römermann, Simon Scheiter, Tim Anders, Sophie Wolf, Mateus Dantas de Paula, Thomas Hickler

## Abstract

Predicting plant functional traits is crucial because traits underpin ecosystem processes. Statistical, machine-learning, and optimality approaches have produced global plant trait predictions. However, the utility of trait-based vegetation models remains unexplored at this scale. We use aDGVM2, an eco-evolutionary trait- and individual-based dynamic global vegetation model (DGVM). aDGVM2 simulates community assembly driven by natural selection. Simulated specific leaf area (SLA) is an emergent outcome of community assembly driven by abiotic conditions and plant competition. We examine (1) how well aDGVM2 simulates global SLA compared to trait data and (2) what drives large differences. Compared to GBIF-derived SLA data, aDGVM2 displays a mean SLA difference of −2.9 m^2^ kg^−1^ (GBIF range ca. 4–35 m^2^ kg^−1^), a root mean square error (RMSE) of 7.26, and a normalised mean absolute error (nMAE) of 26.56 %. Published approaches showed differences of −4.83 to 2.71 (mean), 4.42–6.81 (RMSE), and 11.64 %–25.23 % (nMAE). aDGVM2 mean differences are comparable with published predictions, while RMSEs and nMAEs are higher. Large aDGVM2 mismatches occur where deciduous and evergreen abundances are inaccurate. Correcting abundance mismatches using observed leaf phenology abundances strongly reduces differences (mean: −0.11 to 0.43, RMSE: 5.86–5.91, nMAE: 15.10 %–19.61 %), yielding the smallest mean difference and second-smallest nMAE of all products compared. Thus, an eco-evolutionary, process-based approach can predict global SLA values as well as the best available products once leaf phenological abundances are accurate. Improving the simulation of these abundances is therefore a viable route towards a process-based understanding of the global distribution of SLA and functional trait diversity under contemporary and future climate conditions.

## 1 Introduction

The role of plant traits in understanding and predicting ecosystem function and services has received increasing attention in recent years (Moreno-Martínez et al., 2018; Thonicke et al., 2020). Functional traits, i.e. morphological, physiological, and phenological characteristics that influence plant performance, affect key demographic processes such as growth, survival, and reproduction. Thus, the distribution of plant traits is key to predicting how ecosystems may respond to ongoing environmental changes (Violle et al., 2007; Moreno-Martínez et al., 2018). Traits such as specific leaf area (SLA), leaf nutrient content, and plant height play a direct role in ecosystem processes, including carbon and nutrient cycling, primary productivity, and evapotranspiration (Butler et al., 2017; Madani et al., 2018; Moreno-Martínez et al., 2018; Thonicke et al., 2020; Gomarasca et al., 2023). In particular, SLA has been shown to be a key trait linked to plant function, and its variation can be related to photosynthetic rates, leaf phenology, leaf longevity, and plant strategy (Reich et al., 1991, 1998). Because plant traits mediate interactions between vegetation and biogeochemical cycles, they ultimately shape ecosystem functioning at multiple scales (Lavorel and Garnier, 2002; Violle et al., 2014; Madani et al., 2018). Functional trait diversity has been shown to enhance ecosystem resilience by promoting complementary resource use and stabilizing ecological processes under environmental change (Sakschewski et al., 2016; Thonicke et al., 2020; Billing et al., 2024). Much promise is held in the development of vegetation modelling frameworks to predict the global distribution of plant traits. Such frameworks are essential to improve our understanding of contemporary trait distributions and potential future climate-driven changes in trait composition, and elucidate linkages between functional traits, trait diversity, and ecosystem functions. This challenge, the linking of traits to ecosystem function, has been aptly deemed a holy grail of trait-based ecology (Chacón-Labella et al., 2023).

Billing et al. (2024) recently demonstrated that specific leaf area (SLA) and high within-community diversity in SLA enhances forest biomass across central and eastern Europe via niche complementarity. This study demonstrated that, when forcing the model using RCP2.6 projected climate (a low-warming scenario), higher diversity in SLA increased future biomass storage in the studied forests by up to 18 %. Should relationships between diversity in SLA and ecosystem carbon functioning prove to be globally ubiquitous, then the accurate prediction of SLA variation in vegetation models is crucial to understand future biomass storage and associated carbon dynamics. A requisite first step towards predicting global diversity in SLA, or any other trait, is the accurate prediction of community mean trait values; indeed, mean trait values have shown utility in representing aggregate composition of functional strategies within communities and have been successfully linked to various components of ecosystem functioning (Garnier et al., 2004; Bruelheide et al., 2018; Neyret et al., 2024). Thus, to improve confidence in trait-based vegetation models’ ability to explore past, contemporary, and future linkages between traits, trait diversity, and ecosystem function, an accurate representation of community mean SLA distributions is required.

Over the past decade, numerous methodological advancements have been made to address the pressing need to understand global trait–environment relationships and produce global wall-to-wall trait maps (van Bodegom et al., 2014; Butler et al., 2017; Madani et al., 2018; Moreno-Martínez et al., 2018; Boonman et al., 2020; Dong et al., 2023). Nevertheless, process-based understanding of trait–environment and trait–ecosystem functional relationships is incomplete (Dechant et al., 2024). In the absence of process-based approaches, many of the methods used to predict global trait distributions have necessarily relied on statistical and machine-learning approaches. These empirical methods require trait data sets at large spatial scale to fit and train models (e.g. the TRY plant database; Kattge et al., 2011), various combinations of climate and soil data, and often avail of the distribution of contemporary vegetation patterns (Dechant et al., 2024).

Empirical approaches advance our understanding of global trait–environment relationships and have been shown to reasonably predict the global distribution of traits. However, predictions from these methods are not fully independent of the environmental predictors (Dechant et al., 2024). Thus, despite their predictive performance, statistical and machine learning models are often limited in their utility to disentangle the interactions between environmental and ecological processes that generate observed trait distributions. This is because the successional and temporal dynamics of trait compositions of plant communities are ultimately governed by interactions between the environment, competition, and demographic processes that affect the growth, reproduction, and mortality of plants (Fisher et al., 2018; Iida and Swenson, 2020).

A recent approach, based on the theoretical underpinnings of optimal resource allocation, differs substantially from the above-mentioned empirical methods (Dong et al., 2023). Dong et al. (2023) predict community mean SLA for evergreen and deciduous vegetation, which, based on their theory, are optimal under the given environmental conditions. Thereby, optimality has several components, such as maximising the lifetime leaf net carbon gain and minimizing the cost of maintaining both photosynthetic and water transport capacities. This approach requires climatic data for temperature, vapour pressure deficit, and incident photosynthetically active radiation to generate optimal SLA values for evergreen vegetation. As such, these predictions are independent of plant trait data, which are used only to assess model performance (Dong et al., 2023). In order to generate global maps of plant traits, this methodology requires information on the distribution and relative abundances of evergreen and deciduous plant strategies, which was derived from land cover data sets. The requirement for data that provide the distribution and relative abundances of evergreen and deciduous vegetation to generate community weighted means (CWMs) makes future predictions of traits less certain. Many of the drivers of contemporary leaf phenological distributions are well understood (Givnish, 2002). However, it remains unclear how the distributions of plant strategies will change in the future, since such changes will be governed by feedbacks between changing abiotic conditions, plant demographic processes, changing relative fitnesses, and biogeographic contingency (Moncrieff et al., 2016a, b).

Dynamic global vegetation models (DGVMs) are process-based models that offer a more mechanistic and flexible approach, simulating feedbacks between abiotic conditions, demographic processes, and resource competition, and increasingly incorporate plant hydraulics (Hickler et al., 2004, 2006; Langan et al., 2017; Papastefanou et al., 2024). Some DGVMs include trait diversity and ecological and evolutionary interactions that shape trait distributions and trait diversity (Pavlick et al., 2013; Scheiter et al., 2013; Verheijen et al., 2013; Sakschewski et al., 2015; Langan et al., 2017; Maréchaux and Chave, 2017; Dantas de Paula et al., 2021; Joshi et al., 2022; Rius et al., 2023). Previous studies have successfully employed trait- based DGVMs to predict plant community traits across various spatial scales. For example, Dantas de Paula et al. (2021) used LPJ-GUESS-NTD to investigate how nutrient cycling influences the process of plant community assembly and ecosystem functioning across an elevational gradient in the Andes of southern Ecuador. This study demonstrated how local eco-evolutionary dynamics and abiotic factors, such as nutrient availability, influence the assembly of plant traits in diverse tropical ecosystems. On a broader scale, Thonicke et al. (2020) applied LPJmL-FIT to simulate functional trait diversity across European natural forests along climatic gradients.

Scaling up trait-based DGVM predictions from regional or local scales to the global scale remains a significant challenge. However, understanding global trait distributions and trait diversity is critical for predicting how plant communities will respond to large-scale environmental changes, such as climate change, land-use change, and atmospheric CO_2_ increases (Purves and Pacala, 2008). Trait-based DGVM development has lagged behind empirical methods to predict trait distributions and diversity due to the complexity of ecological systems and the difficulty of parameterising such models at large spatial resolutions. Nonetheless, their ability to link predictions to underlying ecological drivers makes them a valuable tool with the potential to improve our understanding of trait distributions and linkages with ecosystem functioning (Scheiter et al., 2013).

In this study we confront the challenge of global process-based trait predictions. For this, we employ a trait- and individual-based DGVM (aDGVM2). aDGVM2 (v1.0-LL; Langan et al., 2025a) is fully described in Langan et al. (2017, 2025b). aDGVM2 incorporates eco-evolutionary processes that influence the assembly of plant communities and trait distributions. It accounts for both abiotic factors, such as climate and soil conditions, and biotic interactions, including competition for light, water, and space. Trait values and the distribution of evergreen and deciduous leaf phenologies are emergent properties of the natural-selection-driven process of community assembly. A plant’s relative reproductive success in a given environment depends on both its own trait values and those of its competitors.

Thus far, no trait-based vegetation model has been used to compare simulated trait values to observations at a global scale. We aim to test aDGVM2’s ability to simulate the global distribution of SLA. Specifically, we (1) examine how well aDGVM2 can simulate global specific leaf area (SLA) by comparing simulations with GBIF-derived SLA estimates and (2) explore drivers of strong deviations.

## 2 Methods

### 2.1 aDGVM2 model description

aDGVM2 is an individual- and trait-based vegetation model which has been fully described in Langan et al. (2017). In the model, every plant can have a potentially unique set of trait values, the initial trait space is constrained by pre-defined trait ranges (Langan et al., 2017, 2025b), and trade-offs between traits constrain individual plant performance. The model behaves like an iterative optimisation algorithm and thus mimics natural selection. In the model, variability of trait values between individuals in a community is permitted, the set of trait values an individual possesses affects its relative reproductive success, and trait values are heritable. Individuals with poorly performing sets of traits have lower relative reproductive success. This modelled process of natural selection results in a filtering of the community of individuals and their trait values through time. Additionally, individuals are assigned a species label to allow reproductive isolation, which was found to be important for the coexistence of different strategies (Scheiter et al., 2013). Each year individuals reproduce, whereby trait values can be crossed between individuals with the same species label and trait values can mutate with fixed probabilities. This iterative process of reproduction, crossover, and mutation through time allows modelled communities to continuously explore trait space and find trait combinations which may improve relative reproductive success.

### 2.2 Natural selection in aDGVM2: emergent community mean SLA

aDGVM2 includes a simplified version of the cohesion–tension theory of sap ascent in plant hydraulics submodules (see supplements of Langan et al., 2017, 2025b, for a full model description). A key adaptive trait within these submodules which regulates the hydraulic performance of plants is the xylem water potential at 50 % loss of conductance (P50; Fig. 1a). In the model we use an empirically defined set of relationships (Markesteijn et al., 2011) to relate P50 to the conductance of sapwood and leaves, and SLA (Fig. 1b). In the model, low P50 (i.e. high cavitation resistance) results in low leaf and sapwood conductivity and low SLA. High P50 (i.e. low cavitation resistance) results in high leaf and sapwood conductivity and high SLA. Thus, P50 defines a trade-off between safety and transport efficiency. Additionally, SLA defines a trade-off between a plant’s ability to quickly accumulate leaf area and leaf longevity. Here, high SLA promotes faster leaf area accumulation, enhancing photosynthesis while reducing leaf hydraulic resistance. High SLA is also related to lower leaf longevity; thus high-SLA leaves turn over faster than low-SLA leaves (see appendix in Langan et al., 2017, for a full model description). Leaf turnover thus reduces total leaf area and influences photosynthetic carbon gain and leaf resistance (Fig. 1b). By linking xylem cavitation resistance (P50) to the plant hydraulic safety–efficiency spectrum and to the plant economic spectrum governing leaf area accumulation and turnover via SLA, these coupled constraints generate a multidimensional trade-off surface upon which natural selection can act to drive community assembly, trait evolution, and emergent coordination between hydraulic and leaf phenological traits (Langan et al., 2025b).

**Figure 1.**
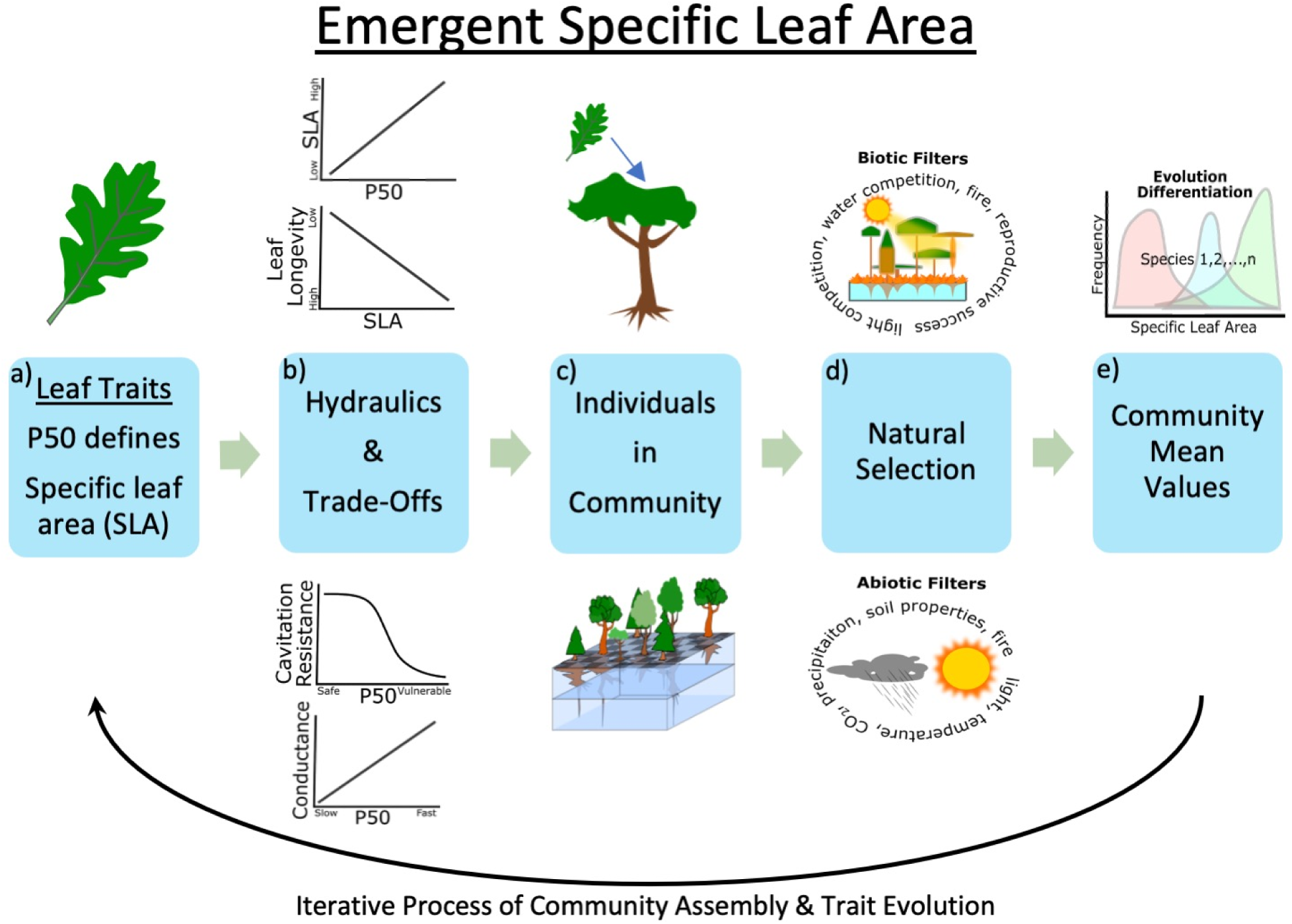
Community assembly in aDGVM2. (a) The value of the adaptive trait, P50 (the water potential at 50 % loss of conductance), defines SLA (see Supplement in Langan et al., 2017, for a full model description). (b) P50 defines a series of trade-offs: (1) high P50 results in high SLA, (2) high SLA results in low leaf longevity, (3) high P50 results in low resistance to cavitation, (4) high P50 results in high conductance. (c) A community of plants is initialised with randomly assigned trait values. (d) Abiotic and biotic filters allow the action of natural selection on plants via reproductive success. (e) The mean traits of communities emerge due to the simulated process of natural selection. Note: lines in (b) are for illustrative purposes only; see appendix in Langan et al. (2017) for a full model description. The figure displays only trade-offs related to selection on SLA.

The model is initialised with a community of plants whose trait values (P50, SLA) are randomly drawn from a uniform distribution with predefined bounds (Fig. 1c). Following initialisation, abiotic (water, light, fire, soil, CO_2_) and biotic (competition for light, water, space, and reproductive success) filters (Fig. 1d) allow natural selection to drive community assembly via relative reproductive success (Fig. 1e). Thus, encoded in the model, through the integration of hydraulic and leaf economic trade-offs, is a hypothesis-driven theory of how natural selection shapes the emergent global distribution of SLA.

### 2.3 Model forcing and simulation setup

aDGVM2 simulates vegetation in 1 ha stands on a daily time step. Therefore, daily climate data are required to force the model. We create time series based on a reference climatology. As reference climatology, we used the Climatic Research Unit (CRU) data set (New et al., 2002), representing climate for the 20th century. This data set provides monthly mean data for temperature, temperature range, precipitation (monthly mean and coefficient of variation), number of rain days, sunshine hours per day, humidity, and number of frost days averaged for the period 1961 to 1990 (that is, 12 monthly values per variable). Soil texture data were obtained from Nachtergaele et al. (2012).

Simulations are run for 900 years for each simulated grid cell using the standard model parameterisation, where the model is initialised with 50 % trees and 50 % grasses to avoid making prior assumptions about initial vegetation distributions (Langan et al., 2017). Fire is turned on; see the model description in Langan et al. (2017) for a description of the fire module. Maximum plant rooting depth is set to 10 m globally. Simulations are run at 1° spatial resolution. We conduct a spin-up of 760 years with CO_2_ concentrations fixed at a pre-industrial (1860) level of 286 ppm to allow vegetation to reach equilibrium. Spinning up the model using pre-industrial CO_2_ concentrations excludes the possibly confounding effects of CO_2_ fertilisation on emergent vegetation. For the last 140 simulation years CO_2_ is increased annually up to 364 ppm, which corresponds to a year 2000 concentration, using recommended annual average pre-2005 global mean concentrations from Meinshausen et al. (2011). For further details on aDGVM2 see the full model description provided as appendices to Langan et al. (2017, 2025b).

### 2.4 Global comparison of aDGVM2 SLA with GBIF data

We initially evaluated the global SLA patterns simulated with aDGVM2 using an observation-based dataset on community-weighted SLA means. This dataset builds on the species-specific SLA observations from the TRY database (Kattge et al., 2011) with spatial species abundance data from GBIF (Wolf et al., 2022; Scheiter et al., 2024). GBIF is a crowd-sourced collection of various datasets, including national surveys and opportunistically sampled individual plant occurrences. Trait products derived from GBIF were found to match closely with sPlot-derived trait data but have better geographic coverage (Wolf et al., 2022; Dechant et al., 2024).

GBIF global SLA maps were derived by integrating GBIF species observations with SLA observations from the TRY database (Kattge et al., 2020), following Wolf et al. (2022) and Scheiter et al. (2024). A total of 257 357 303 species observations were downloaded from GBIF on 2 June 2023 (GBIF.org, 2023). This dataset was pre-filtered for “Observation”, “Human observation”, or “Occurrence” records, ensuring no geospatial issues, no related records, a minimum distance of 1500 m from country centroids, and occurrence status set to “present” (since GBIF includes true absence data). Additional filtering via “CoordinateCleaner” (Zizka et al., 2019) removed records with > 10 km coordinate uncertainty, > 0.1° precision, ocean locations, and common errors (e.g. equator/meridian artefacts), retaining only species-level identifications. The cleaned observations were then linked to TRY’s gap-filled dataset via species names (Schrodt et al., 2015), yielding 192 667 225 observations. Overall, 90 % of GBIF observations and 24 % of GBIF species were matched to 70 % of TRY species. To reduce computational load and redundancy, we spatially subsampled observations into 2591 km^2^ equal-area hexagons (“dggridR”; Barnes and Sahr, 2023), approximating 0.5° at the Equator. Up to 10 000 observations were retained per hexagon, resulting in 31 808 221 observations. These were aggregated using a mean function into a 1° global raster grid. For the GBIF-based approach, SLA maps were created for three plant functional type (PFT) combinations: (1) grasses and herbs (grasses), (2) trees and shrubs (woody plants), and (3) all PFTs combined. Species were assigned to PFTs via a majority vote based on TRY’s PFT data (trait ID 197). The creation of separate functional types facilitates comparisons between functional groups and with PFTs in vegetation models. For the trait comparisons presented here, SLA maps for tree and shrub PFTs (2 above) were compared to aDGVM2 CWM SLA for woody plants.

### 2.5 Statistical comparisons

Although the use of correlative goodness-of-fit metrics to evaluate model performance against observations is pervasive, such correlative methods, while familiar, can be highly sensitive to extreme values. Correlative methods are insensitive to additive and proportional differences, whereby a high coefficient of determination (*r*^2^) can be achieved even when there are large differences between simulated/predicted and observed values (Legates and McCabe, 1999). A trivial example would be that a perfect linear relationship could be achieved if a model consistently predicted SLA to be an order of magnitude larger than observations. For these reasons, we assess model performance using a series of metrics which focus on the magnitudes of differences, as recommended for process-based model evaluation (Legates and McCabe, 1999).

To evaluate the performance of aDGVM2 SLA predictions and statistical, machine learning, and optimality-based SLA predictions against observational datasets, we compare mean differences, calculate 95 % confidence intervals about mean differences, root mean square error (RMSE), and normalised mean absolute error (nMAE). Mean differences provide a straightforward measure of bias and allow examination of whether the model consistently overestimates or underestimates SLA. Confidence intervals provide an intuitive method to assess the potential range of mean differences. RMSE provides an unbounded measure of error which is sensitive to inflation due to large differences; nMAE provides error estimates which are less sensitive to large differences, while large differences between RMSE and nMAE are indicative of the presence of large differences (outliers) between simulated/predicted and observed SLA (Legates and McCabe, 1999).

### 2.6 Assessing SLA differences across terrestrial ecoregions

To gain further insight into differences between simulated SLA and observed values, the WWF terrestrial ecoregions data (Olson et al., 2001) are used to delimit differences on a per-biome basis.

### 2.7 Gap filling and re-weighting SLA values

#### 2.7.1 Gap filling SLA values

aDGVM2 simulations do not always predict deciduous and evergreen vegetation in all grid cells where these leaf phenology types are observed. Data on the relative abundances of evergreen and deciduous trees and shrubs from the ESA Climate Change Initiative (ESA-CCI; Hartley et al., 2017) for the year 2000 were downloaded and aggregated to the same 1° resolution as model simulations. ESA-CCI data are initially used to identify simulated grid cells where either leaf phenological type is observed but not simulated.

Three different methods are applied to fill SLA values of deciduous and evergreen vegetation in grid cells where they are observed but not simulated. The following methods are applied separately for deciduous and evergreen leaf phenological types: (1) the global mean values of simulated SLA are used to fill gaps, (2) WWF terrestrial ecoregion data (Olson et al., 2001) are used to return the mean value of simulated SLA for each of the world’s biomes, and (3) WWF data are again used to return the mean value of simulated SLA for each biome in each biogeographical realm (Olson et al., 2001).

This process produces three gap-filled SLA rasters for deciduous and evergreen vegetation, respectively:

1. SLA filled using global mean values;
2. SLA filled using biome-specific mean values;
3. SLA filled using biome- and realm-specific mean values.

#### 2.7.2 Re-weighting global SLA values

The ESA-CCI data on the relative abundances of evergreen and deciduous trees and shrubs are then used to re-weight gap-filled SLA rasters. Abundances were divided by the total abundance of evergreen and deciduous trees and shrubs to ensure abundances summed to one. Adjusted SLA values are calculated for deciduous and evergreen leaf phenological types by multiplying each raster with the respective abundance and then adding the SLA values of adjusted deciduous and evergreen rasters together as follows:

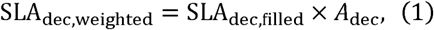

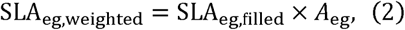

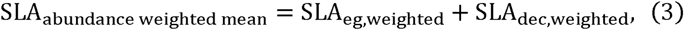

where *A*_dec_ and *A*_eg_ are the normalised ESA-CCI abundances of deciduous and evergreen vegetation, respectively.

### 2.8 Comparing aDGVM2 simulated, empirical, and theory-derived SLA with observations globally

Empirical and theory-derived SLA datasets were downloaded from the supplement of Dong et al. (2023) at 0.5° resolution and aggregated to 1°. These data include empirically derived global trait predictions (van Bodegom et al., 2014; Butler et al., 2017; Madani et al., 2018; Moreno-Martínez et al., 2018; Boonman et al., 2020). Hereafter, each dataset used to contextualise aDGVM2 performance is referred to by the name of the first author.

In this paper it is not our intent to provide detailed descriptions of the methods used to derive the empirical predictions which we use for comparison; rather, we compare aDGVM2 simulated SLA with published products to assess relative performance. For a detailed discussion of the various products used for comparison see the respective papers; a detailed comparison of all empirically derived products is provided by Dechant et al. (2024).

## 3 Results

### 3.1 Global differences between simulated and observation-based SLA

Comparing global differences of aDGVM2 simulated SLA with GBIF-derived trait maps reveals many areas of good agreement, where differences between simulated and observed SLA were small (Fig. 2a–c). Also apparent in the global maps are large areas of bias where SLA is underestimated by aDGVM2. Areas of bias where SLA is strongly overestimated are generally restricted to high-northern-latitude boreal and tundra areas.

**Figure 2.**
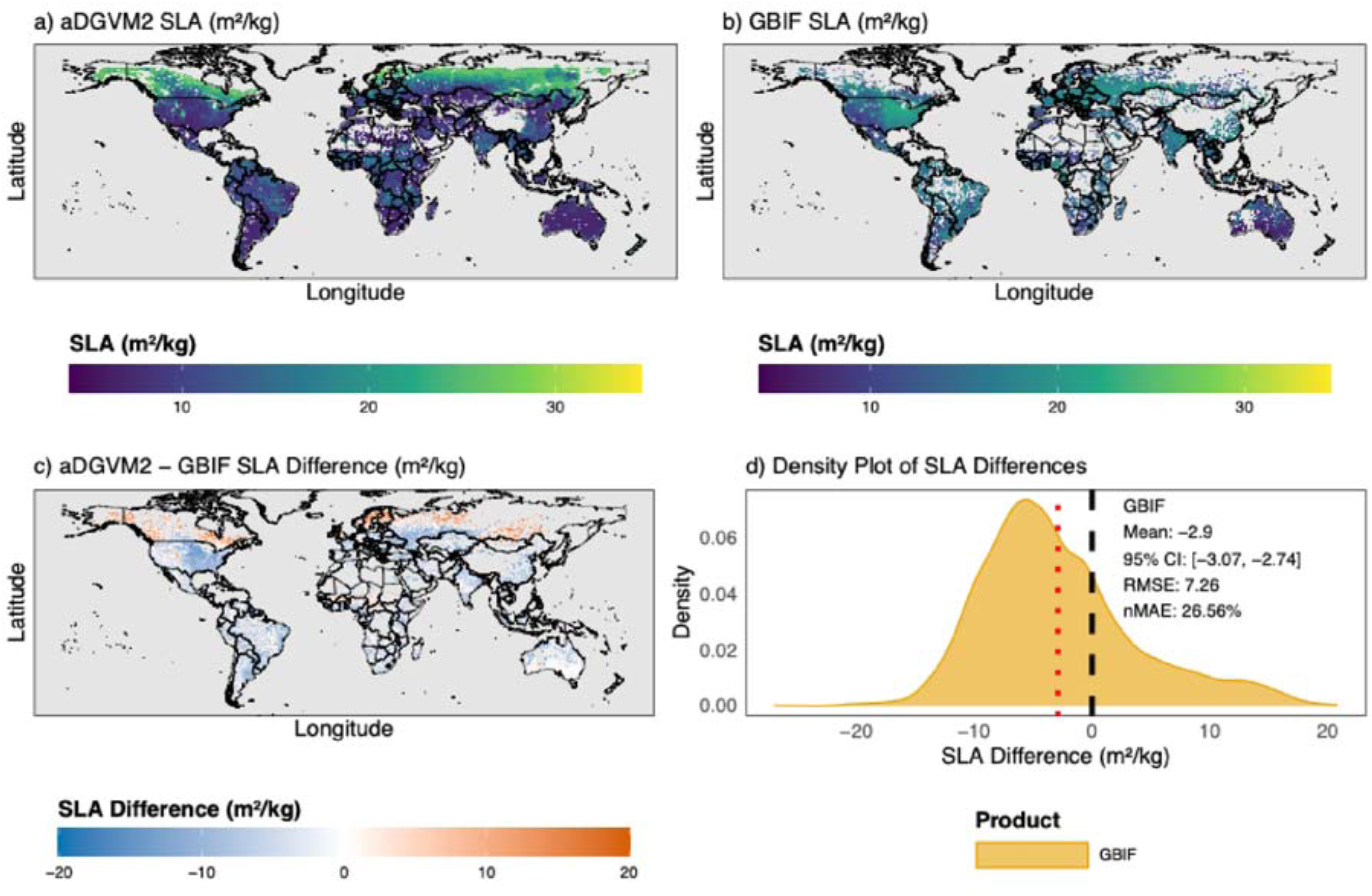
aDGVM2 simulated vs. GBIF-derived SLA. (a) Spatial distribution of aDGVM2 SLA. (b) Spatial distribution of GBIF-derived SLA. (c) Spatial distribution of differences (aDGVM2 − GBIF). (d) Density plots of aDGVM2 − GBIF SLA differences. The red vertical line indicates the mean difference.

Global comparisons reveal that aDGVM2 simulated SLA differs from GBIF data by an average of −2.9 m^2^ kg^−1^ (Fig. 2d) and thus displays a general global bias towards underestimation. A 95 % confidence interval for the difference in the means ranges between −3.07 and −2.74. This indicates statistically significant mean differences between simulated and observed SLA. The root mean square error of differences (RMSE) is 7.26, while the normalised mean absolute error (nMAE) is 26.56%.

### 3.2 Biome-specific differences between simulated and observed SLA

To better understand the differences between simulated and observed SLA, differences are compared at the biome level (Fig. 3). This analysis reveals the same general patterns of bias observed in the global patterns. In the boreal forests/taiga (Fig. 3g) and tundra (Fig. 3l) biomes SLA is systematically overestimated. These biomes also exhibit the lowest data coverage (38.6 %, 39.5 %), i.e. the percentage of simulated grid cells for which GBIF-derived SLA was available. The vast majority of comparisons in other biomes reveal systematic underestimation of SLA (Fig. 3). The largest SLA underestimations are in flooded grasslands and savannas (Fig. 3j; −5.56; data coverage: 53.33 %), tropical and subtropical moist broadleaf forests (Fig. 3b; −5.06; data coverage: 65.71 %), and temperate broadleaf and mixed forests (Fig. 3e; −4.72; data coverage: 86.62%); comparisons for these biomes also exhibit the highest RMSE and nMAE.

**Figure 3.**
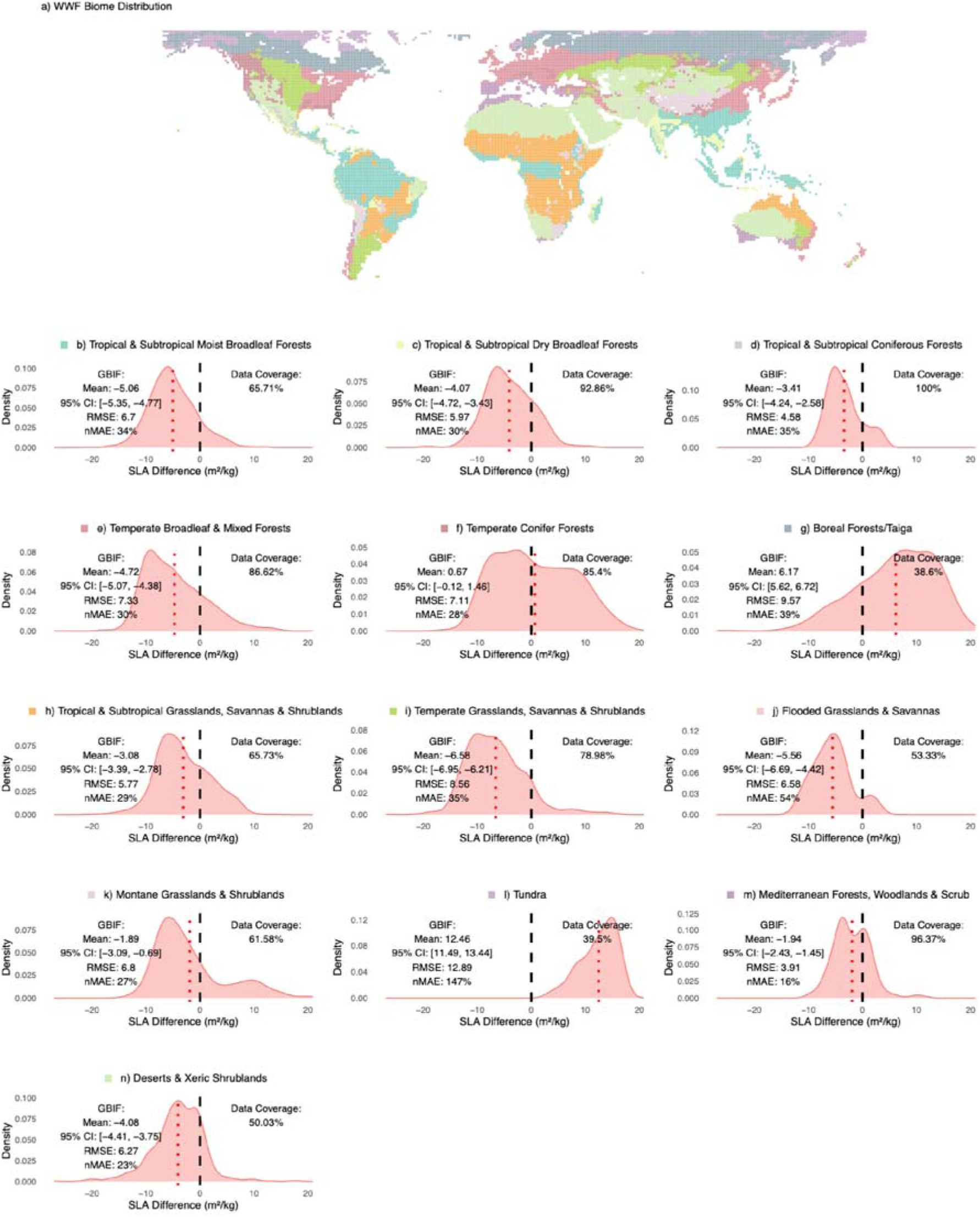
Per-biome differences between aDGVM2 simulated SLA and GBIF-derived SLA. (a) Map of the WWF biome distribution. (b–n) Biomes.

For temperate coniferous forests (Fig. 3f) and montane grasslands and shrublands (Fig. 3k) differences between simulated and observed SLA are smaller. The mean difference between simulated SLA and GBIF-derived SLA in temperate coniferous forests is 0.67 with a 95 % confidence interval of (−0.12, 1.46) and a data coverage of 85.4 %. The mean difference between simulated SLA and GBIF-derived SLA in montane grasslands and shrublands is −1.89 with a 95 % confidence interval of (−3.09, −0.69) and a data coverage of 61.58 %.

### 3.3 Potential sources of error: emergent differences between aDGVM2 simulated SLA for evergreen and deciduous leaf phenologies

Optimality-theory-derived SLA (Dong et al., 2023) differs for deciduous and evergreen vegetation, and SLA comparisons between predicted and observed values were carried out for evergreen and deciduous taxa separately and also by weighting the relative abundances of evergreen and deciduous leaf phenologies. Indeed, linkages between SLA and photosynthesis have been shown to differ between evergreen and deciduous vegetation (Reich et al., 1998; Kröber et al., 2014). Such differences between evergreen and deciduous vegetation suggest two potential sources of error for the systematic underestimation of SLA across many biomes in aDGVM2: (1) observed differences between evergreen and deciduous SLA may not be captured by aDGVM2, and (2) the relative abundances of simulated evergreen and deciduous vegetation may be incorrect.

The separate examination of the global distribution of simulated SLA for deciduous and evergreen vegetation (Fig. A1a, b) reveals that each leaf phenological type displays a large amount of variability in mean community values. Evergreen SLA displays a global range of ca. 7–27 m^2^ kg^−1^ (Fig. A1c), has a global mean of 12.08 m^2^ kg^−1^, and displays relatively lower values in dry regions, while the highest values are simulated in the boreal forests and tundra of northern latitudes. Deciduous vegetation displays a global range of ca. 7–35 m^2^ kg^−1^ (Fig. A1c), has a global mean of 20.68 m^2^ kg^−1^, and displays relatively lower values in savannas, while higher values are simulated in tropical forests and the temperate forests of northern latitudes. While evergreen and deciduous leaf phenologies display clear differences in mean values (Fig. A1c), large geographical areas where no deciduous vegetation is simulated are apparent from our simulations (Fig. A1b). Taken together, these results show that simulated deciduous and evergreen SLA do indeed differ. However, they additionally highlight that the simulated absence of deciduous vegetation across large geographical areas and potential mismatches in the relative abundances of evergreen and deciduous leaf habits may be responsible for the systematic underestimation of SLA.

### 3.4 Potential sources of error: simulated vs. observed abundance of evergreen and deciduous leaf phenologies

To assess whether systematic SLA underestimations are a result of potential errors associated with the incorrect simulation of the relative abundances of evergreen and deciduous vegetation, we compare our simulated abundances with satellite-derived abundances (Fig. A2). This comparison reveals that aDGVM2 simulated abundances differ substantially from satellite-derived data across large spatial scales (Fig. A2a, b).

While there are many areas of good agreement, particularly across the tropics, at the global scale the simulated abundance of evergreen vegetation is systematically overestimated (Fig. A2a). Correspondingly, the abundance of deciduous vegetation is systematically underestimated (Fig. A2b). These results indicate that the simulated abundance of evergreen vegetation is about 30 % too high (Fig. A2c).

### 3.5 Biome-specific SLA changes between original vs. gap-filled and abundance-adjusted SLA values

We then explicitly examine whether the systematic underestimation of simulated SLA is the result of abundance differences between simulated and observed abundances of evergreen and deciduous phenologies. Our analysis revealed that, of the 41 261 simulated grid cells, there were 3065 gaps where deciduous vegetation is observed but not simulated and 259 gaps where evergreen vegetation is observed but not simulated. Comparing aDGVM2 gap-filled and abundance-adjusted SLA (see Sect. 2.7.1 and 2.7.2) with GBIF-derived trait maps reveals that, in biomes where SLA is systematically too low, adjusted predictions display higher mean SLA across all of these biomes (Fig. 4).

**Figure 4.**
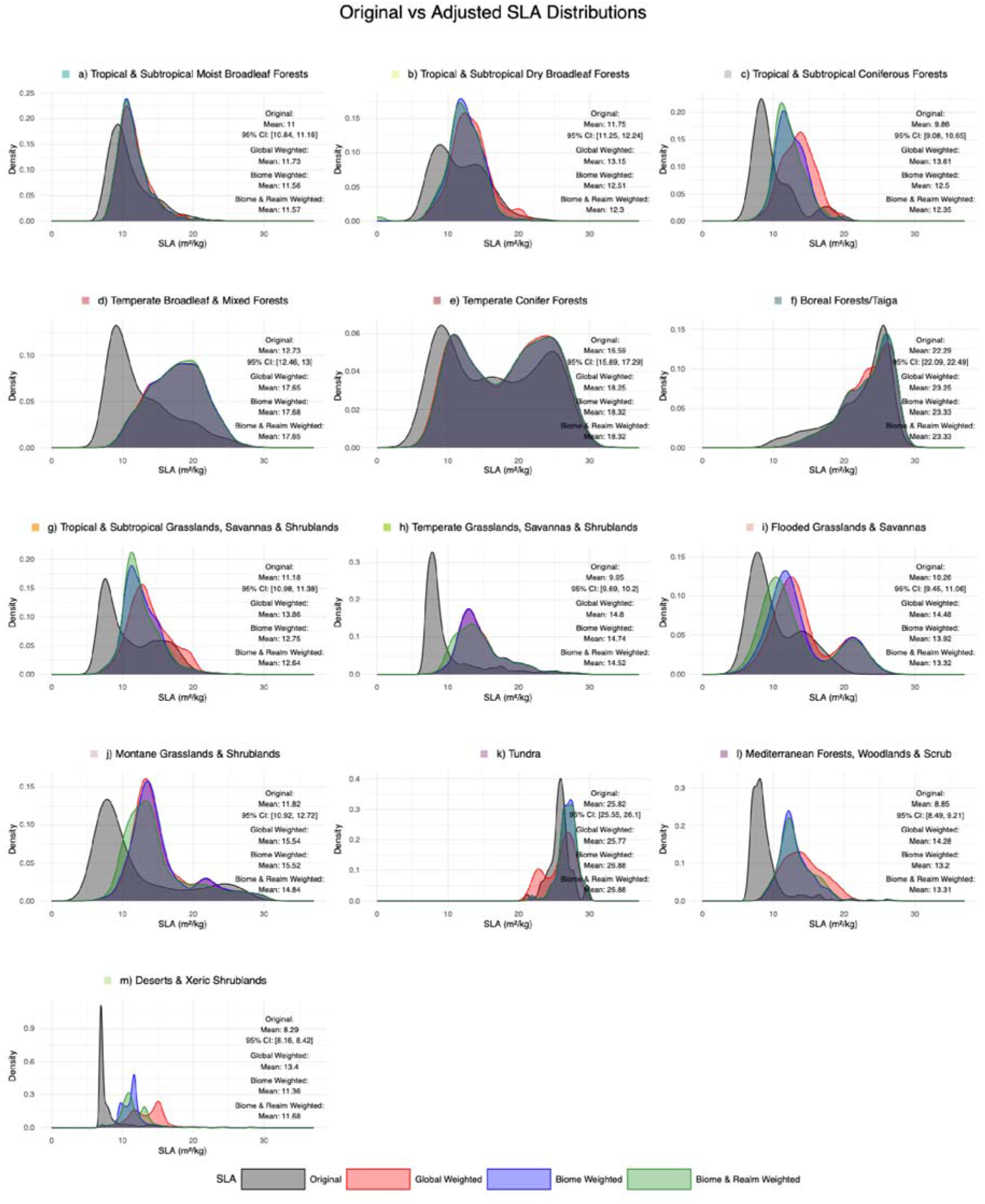
Per-biome density plots of original and adjusted aDGVM2 SLA values.

In the boreal forest and tundra biomes, where unadjusted SLA is overestimated, adjusted SLA does not differ much from the unadjusted SLA (Fig. 4f, k). The smallest increases are found in tropical and subtropical moist broadleaf forests (Fig. 4a; increases of 0.56–0.73), tropical and subtropical dry broadleaf forests (Fig. 4b; increases of 0.55–1.4), and temperate conifer forests (Fig. 4e; increases of 1.66–1.73).

In all other biomes (Fig. 4), increases in SLA are higher. The largest increases in SLA are found for temperate broadleaf and mixed forests (Fig. 4d), which is the biome with the biggest mismatch between the simulated abundances of deciduous and evergreen leaf phenologies and between unadjusted and observed SLA (Fig. 3e), with an increase in SLA of 4.92–4.95.

### 3.6 Global comparisons of aDGVM2 simulated, empirically predicted, and theory-derived SLA

In order to better evaluate the performance of aDGVM2 unadjusted and adjusted SLA, we compare our simulated output with GBIF-derived SLA data and include in our analysis additional comparisons of SLA predictions from a range of previously published products. This approach allows us to better assess whether aDGVM2 SLA predictions are comparable to those of published products, assess the effectiveness of adjustments, and identify areas of bias.

#### 3.6.1 Comparing biases in SLA in unadjusted vs. adjusted aDGVM2 predictions

Comparing unadjusted and adjusted aDGVM2 SLA globally with GBIF-derived trait distributions reveals that all of the adjustment methods strongly reduced global biases in simulated SLA. The different adjustment methods all reduce global mean differences, RMSE, and nMAE (Figs. 5 and 6a–d; Table 1).

**Figure 5.**
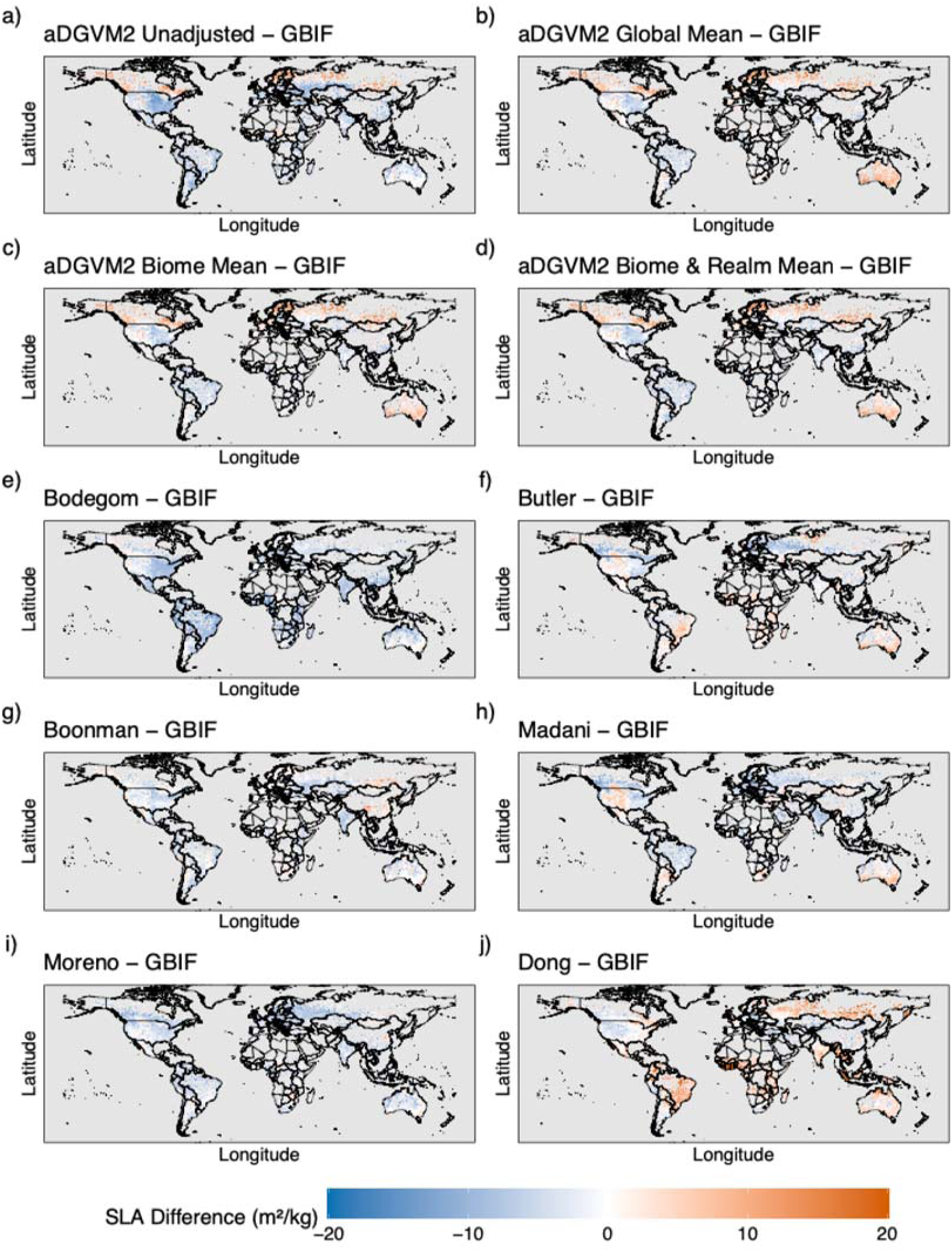
Spatial distributions of global SLA differences between aDGVM2 simulations, published maps, and GBIF-derived SLA. (a) aDGVM2 Unadjusted, (b) aDGVM2 Global Mean, (c) aDGVM2 Biome Mean, (d) aDGVM2 Biome and Realm Mean, (e) van Bodegom et al. (2014), (f) Butler et al. (2017), (g) Boonman et al. (2020), (h) Madani et al. (2018), (i) Moreno-Martínez et al. (2018), (j) Dong et al. (2023).

**Figure 6.**
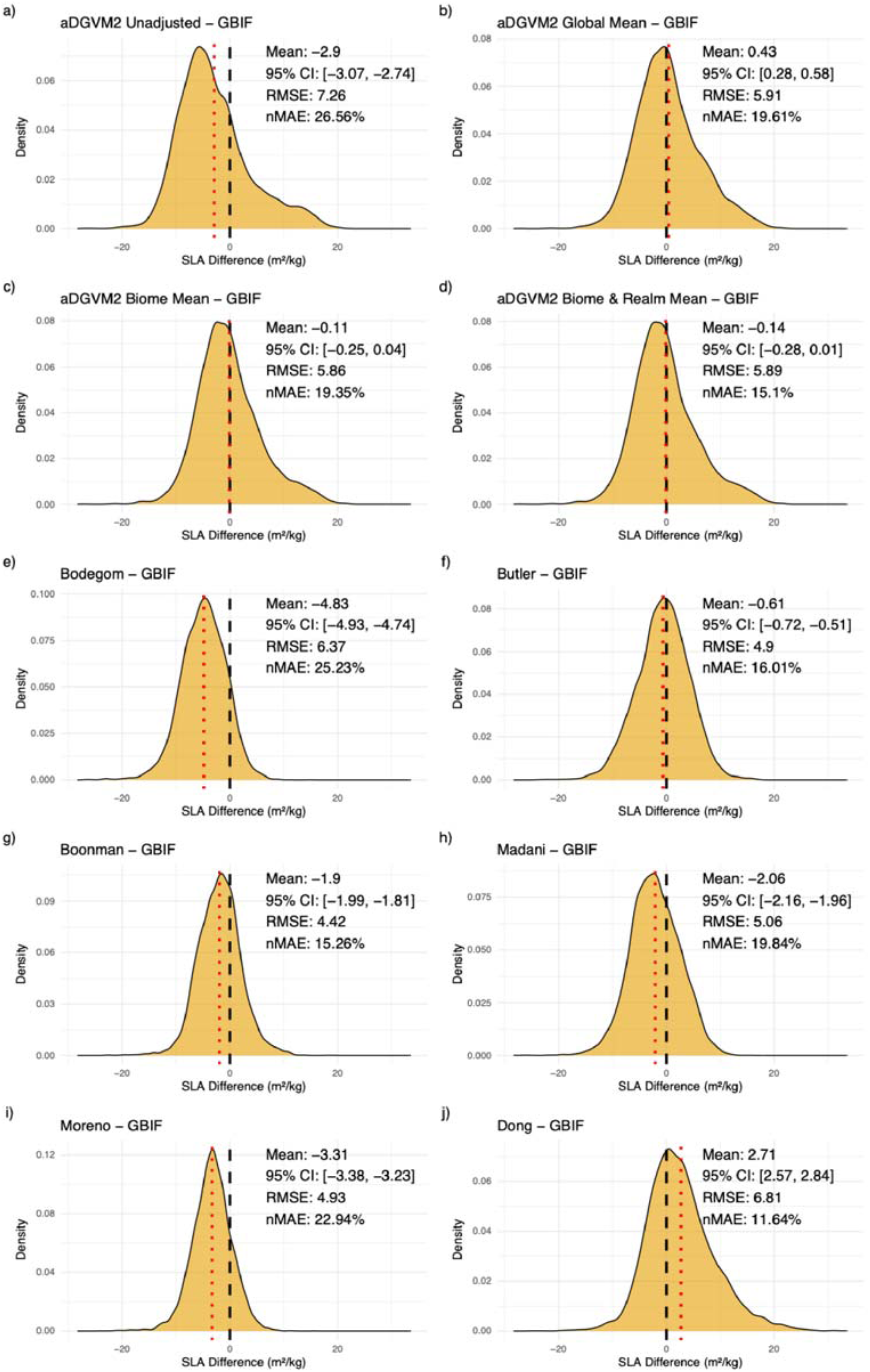
Density plots of global SLA differences between aDGVM2 simulations, published maps, and GBIF-derived SLA data. (a) aDGVM2 Unadjusted, (b) aDGVM2 Global Mean, (c) aDGVM2 Biome Mean, (d) aDGVM2 Biome and Realm Mean, (e) van Bodegom et al. (2014), (f) Butler et al. (2017), (g) Boonman et al. (2020), (h) Madani et al. (2018), (i) Moreno-Martínez et al. (2018), (j) Dong et al. (2023).

**Table 1.** Mean differences between unadjusted and adjusted aDGVM2 simulated SLA, a suite of previously published SLA predictions, and GBIF-derived trait data. Shown are mean differences, a 95 % confidence interval (CI) for the differences in means, the root mean square error (RMSE), and the normalised mean absolute error (nMAE) in percent.

| SLA differences | Mean difference | CI lower | CI upper | RMSE | nMAE (%) |
| --- | --- | --- | --- | --- | --- |
| aDGVM2 Unadjusted – GBIF | –2.90 | –3.07 | –2.74 | 7.26 | 26.56 |
| aDGVM2 Global Mean – GBIF | 0.43 | 0.28 | 0.58 | 5.91 | 19.61 |
| aDGVM2 Biome Mean – GBIF | –0.11 | –0.25 | 0.04 | 5.86 | 19.35 |
| aDGVM2 Biome & Realm Mean – GBIF | –0.14 | –0.28 | 0.01 | 5.89 | 15.10 |
| Bodegom – GBIF | –4.83 | –4.93 | –4.74 | 6.37 | 25.23 |
| Butler – GBIF | –0.61 | –0.72 | –0.51 | 4.90 | 16.01 |
| Boonman – GBIF | –1.90 | –1.99 | –1.81 | 4.42 | 15.26 |
| Madani – GBIF | –2.06 | –2.16 | –1.96 | 5.06 | 19.84 |
| Moreno – GBIF | –3.31 | –3.38 | –3.23 | 4.93 | 22.94 |
| Dong – GBIF | 2.71 | 2.57 | 2.84 | 6.81 | 11.64 |

Comparisons with GBIF-derived SLA show that the 95 % confidence intervals for the difference in the means for aDGVM2 Biome Mean (Fig. 6c; Table 1) and aDGVM2 Biome and Realm Mean (Fig. 6d; Table 1) both include zero. Overestimation biases remain in the boreal forest and tundra biomes (Fig. 5a–d). The adjustment method introduced an overestimation bias in Australia. All other global SLA underestimations are strongly reduced.

#### 3.6.2 Comparing aDGVM2 performance against published SLA predictions

Comparing differences between aDGVM2 unadjusted SLA and GBIF data with published products reveals that the mean differences, RMSE, and nMAE of unadjusted predictions represent an outlier when compared to published products. Unadjusted differences show the third-highest mean differences across products (Table 1), the highest RMSE, and the highest nMAE.

Global mean, biome mean, and biome and realm mean adjustments all reduce mean differences between aDGVM2 and GBIF-derived SLA (Fig. 6; Table 1). Biome mean adjusted values display the lowest mean difference between predicted and GBIF-derived SLA of all products (Fig. 6; Table 1). Biome and realm mean differences are marginally more negative than biome mean adjusted values and display the second-lowest mean differences across products; adjusted predictions display an RMSE comparable to those of other products and have the second-lowest nMAE.

There are a number of spatial similarities and dissimilarities between aDGVM2 SLA and that of other products. Both aDGVM2 and Dong et al. (2023) overestimate SLA in boreal and tundra biomes (Fig. 5). In Australia, adjusted aDGVM2, Butler et al. (2017), Madani et al. (2018), and Dong et al. (2023) data tend to overestimate SLA (Fig. 5); this overestimation is not present in unadjusted aDGVM2 SLA predictions. Various ranges of underestimation of SLA in the temperate broadleaf and mixed forests of North America and tropical forests are common for aDGVM2, van Bodegom et al. (2014), Madani et al. (2018), Moreno-Martínez et al. (2018), and Boonman et al. (2020).

## 4 Discussion

### 4.1 Global SLA distribution and major areas of mismatch

Initial comparisons of unadjusted aDGVM2 simulated SLA reveal two major biases when compared to GBIF-derived trait maps. Apparent from comparisons were overestimations of SLA in northern-latitude tundra and boreal forest biomes and pervasive underestimation of SLA in temperate biomes (Fig. 2).

#### 4.1.1 Areas of SLA overestimation

The overestimation of SLA in tundra and boreal forest biomes is not unexpected, as aDGVM2 does not yet include a number of the environmental drivers of low SLA in these regions; we note, however, that these biomes exhibited the lowest data coverage of all biomes. Givnish (2002) includes the dominance of low-SLA evergreen vegetation, rather than higher-SLA deciduous vegetation, in highly seasonal boreal forests as one of the three great paradoxes that no model could reasonably explain. Givnish suggests interactions between infertile soils, high levels of leaching, and low decomposition rates in boreal regions as potential explanations for evergreen dominance in these areas. Indeed, Shaver et al. (2001) found that 15 years of nitrogen and phosphorus fertilisation of Alaskan tundra resulted in a large shift in vegetation composition from communities dominated by graminoid, evergreen, deciduous, and moss species to communities strongly dominated by deciduous shrubs. aDGVM2 does not currently include nutrient dynamics and cycling; thus, it is unsurprising that the model predicts higher SLA than GBIF-derived SLA data and a high abundance of deciduousness compared to ESA-CCI estimates. Further, Anderson et al. (2020) showed that in boreal forests, SLA increases with increasing permafrost active layer depth, i.e. the maximum depth of soil available to plants above permafrost. Indeed, Sakschewski et al. (2021) convincingly demonstrate the importance of variable plant rooting and soil depth in mediating the distribution of evergreen vegetation in tropical South America. These findings echo previous findings with aDGVM2 which demonstrate that changes in soil depth can increase or decrease P50, and thus SLA, and the abundance of evergreen and deciduous trees across a precipitation gradient (Langan et al., 2017). As the current simulations are a first global test of the model’s ability to predict SLA, we set the global soil depth to a maximum of 10 m. This setup will provide a valuable initial status quo of model performance, as it will allow us to compare future simulations with variable soil depths to this 10 m baseline.

#### 4.1.2 Areas of SLA underestimation: leaf phenology and re-weighting community mean SLA

In contrast to the SLA overestimation in tundra and boreal forest areas, underestimation of SLA dominates the remainder of the globe (Fig. 2). These underestimations are particularly striking in temperate broadleaf and mixed forest biomes. This obvious discrepancy suggests that the mix of leaf phenological habits in this, and potentially all, biomes may be inaccurately simulated. Further, Dong et al. (2023) highlight the different predictions of SLA for different leaf phenological habits. We thus investigate whether aDGVM2 simulated SLA differed for evergreen and deciduous trees (Fig. A1). This analysis confirmed that, while there is considerable overlap in simulated SLA, deciduous and evergreen vegetation tend to emerge from the simulated process of community assembly with very different SLA distributions and mean values, with evergreen vegetation tending to have lower SLA (global mean = 12.08 m^2^ kg^−1^) in comparison to deciduous vegetation with higher SLA (global mean = 20.68 m^2^ kg^−1^). Together these results suggest that, while SLAs for evergreen and deciduous vegetation generally differ in aDGVM2, mismatches in the relative abundances of these leaf phenological habits may explain the areas of global SLA underestimation.

In order to ascertain whether mismatches in the simulated relative abundances of evergreen and deciduous leaf habits are responsible for SLA underestimation, we compare abundances to ESA-CCI abundances (Fig. A2). This analysis reveals that, while the relative abundances were reasonably well simulated across the tropics, a focal area of initial model development and testing, across the rest of the globe deciduous vegetation is approximately 30 % more abundant than simulated by aDGVM2.

The almost globally ubiquitous overestimation of evergreen abundance (underestimation of deciduous abundance) provides further evidence that SLA mismatches may not be due to incorrect simulation of SLA for evergreen and deciduous vegetation but rather the result of inaccuracies in simulating evergreen and deciduous relative abundances. Recalculating the community mean simulated SLA by re-weighting based on observed relative abundances (Fig. 4) shows that adjustments to the relative abundances of evergreen and deciduous leaf phenologies strongly increase the mean SLA in almost all biomes (Fig. 4).

### 4.2 Comparisons between aDGVM2 simulated SLA and published products

To situate our simulation results for predicted SLA within the global context, we compare them to a series of previously published products. These comparisons reveal that, while unadjusted SLA predictions tend to perform towards the poorer end of the spectrum, adjusted SLA predictions are among the best of all previously published products in terms of minimal mean differences, low RMSE, and low nMAE when compared to GBIF-derived SLA data. These results strongly suggest that a process-based understanding of trait-based community assembly, which incorporates demographic processes, i.e. the birth, growth, and mortality of individual plants, is plausible, and they highlight the need to improve representations of the drivers of evergreen and deciduous vegetation in aDGVM2.

### 4.3 The surprising convergence of aDGVM2-based and optimality-theory-based predictions of SLA

While the majority of previously published products that predict global SLA were statistical or empirical in nature, Dong et al. (2023) present a theory-driven set of hypotheses to predict global SLA. Their approach estimates optimal SLA for deciduous and evergreen vegetation separately. As input, this approach requires abiotic data, leaf area index, and the ratio of actual to potential evapotranspiration. Community-weighted mean values are then calculated using observed abundances of evergreen and deciduous vegetation. Despite strong differences between this approach and that of aDGVM2, both produce remarkably similar performance.

In aDGVM2, plant hydraulic constraints, trade-offs, and linkages between SLA and leaf longevity (a key optimisation assumption in Dong et al., 2023), in combination with abiotic conditions and competition between plants, allow natural selection to drive the process of community assembly and produce community mean SLA values. This model structure represents a formally coded theory of how natural selection drives the process of community assembly across broad abiotic and biotic gradients.

The similar performance of these two separate theories in representing the global distribution of SLA is remarkable. Dong et al.’s (2023) parameter-sparse empirical formulation of optimum specific leaf area focuses on the leaf economics spectrum, while aDGVM2’s natural-selection-driven formulation focuses more on plant hydraulics, trade-offs constraining plant performance, competition, and an explicit representation of plant demographic processes. The convergence of results from these two theories suggests that integrated representations of the leaf economics spectrum, as presented by Dong et al. (2023), and of plant hydraulics may provide a plausible route towards a mechanistic and process-based understanding of the biogeography of plant traits (Kröber et al., 2014; Xu et al., 2021).

### 4.4 Future directions

The drivers of SLA, and those influencing leaf phenology and the coexistence, relative abundance, or competitive exclusion of evergreen and deciduous trees, interact. Some of the key factors driving these distributions are already included in aDGVM2, e.g. temperature, water availability in interaction with plant hydraulics, fire disturbance, and the volume/depth of the rooting zone. Yet, the exact nature of how these factors interact to influence the process of community assembly remains poorly understood, in reality and in the model. Soil fertility is another important driver of SLA (Reich et al., 1997, 1998) and leaf phenology (Givnish, 2002), particularly in the nutrient-limited tundra and boreal forests where the model systematically overestimates SLA. Thus, the inclusion of nutrient dynamics within aDGVM2 represents a plausible path to improve the simulated SLA distribution and relative abundances of evergreen and deciduous vegetation.

This paper presents the first global aDGVM2 simulations; our aims were to assess model performance and provide a benchmark which can be used to compare with future iterations. As such, we did not investigate a number of the known drivers of SLA and leaf phenology. Soil depth is known to affect the SLA of plant communities (Anderson et al., 2020), while pedological conditions can result in atypical vegetation across vast areas of the globe (Walter and Breckle, 2002). It has previously been shown with aDGVM2 (Langan et al., 2017) that soil depth affects the traits (e.g. SLA/P50 and the proportion of deciduous vegetation) in a community.

Additionally, fire, acting as a global herbivore (Bond and Keeley, 2005), can significantly alter the distribution of simulated vegetation in comparison to simulations where fire was turned off (Bond et al., 2005). For example, Bond et al. (2005) showed that, using the SDGVM vegetation model, the presence or absence of fire significantly altered the global distribution of dominant vegetation types. This study was the first to demonstrate such broad-scale effects of fire on the distribution of vegetation. However, at the time, the enormous diversity between plants (Kattge et al., 2011) was aggregated into four functional groups (C_3_ grasses, C_4_ grasses, angiosperms, and gymnosperms). Given that fire occurs frequently across four of the world’s major biomes (temperate and tropical grasslands and savannas, Mediterranean shrublands, and boreal forests) and can drive alternative vegetation states (Pausas et al., 2017), utilisation of the higher-resolution representation of functional trait diversity within aDGVM2 to explore the extent to which fire might mediate the process of community assembly is a logical next step.

## 5 Conclusions

The results presented in this study highlight the potential of process-, trait-, and individual-based vegetation models which employ an eco-evolutionary approach to predict the outcome of community assembly. Previously, such an approach was thought to be impossible (Shipley, 2009). To paraphrase Shipley’s reservations, developing a population-based model where the outcome of simulations would be community mean trait values is similar to an attempt to predict the behaviour of a gas by modelling the dynamics of the individual molecules colliding through time. Yet, the presented model, which simulates populations of plants, where the initial trait distributions are randomly initialised and natural selection drives the process of community assembly and trait evolution, clearly demonstrates an ability to predict the global patterns of SLA as well as previously published approaches. Such a representation of community assembly embedded within a vegetation model, while complex, allows the examination of linkages between traits and ecosystem function, as these are explicitly modelled. Further, the presented approach has demonstrated utility in predicting mean traits; however, the model does not predict a single mean value of SLA for a particular combination of environmental conditions. Rather, the model predicts the co-occurrence of traits and plant strategies, i.e. the functional diversity across the globe, which is thought to underpin ecosystem resistance and resilience. The model can also be used to simulate changes in functional diversity through time and examine interactions between how changes in climate may affect trait distributions and how changes in trait distributions may affect ecosystem function, which can feed back on atmospheric processes. Thus, this approach has the potential to answer critically important aims of functional ecology by linking trait distributions to ecosystem function globally (Chacón-Labella et al., 2023).

## Code availability

The aDGVM2 model code (v1.0-LL) is available on Zenodo at https://doi.org/10.5281/zenodo.16265491 (Langan et al., 2025a). The R scripts used to produce all data figures and Table 1 are available, together with the simulation output, at https://doi.org/10.5281/zenodo.23768446.

## Data availability

aDGVM2 simulation output (community mean SLA for all, evergreen, and deciduous vegetation; trait pool; community abundances) and the derived abundance-weighted SLA rasters are available at https://doi.org/10.5281/zenodo.23768446. Third-party data used in this study are not redistributed but are available from their original sources: GBIF-derived trait maps (Wolf et al., 2022) at https://github.com/tejakattenborn/GBIF_trait_maps, based on the GBIF occurrence download (GBIF.org, 2023); ESA-CCI land cover data (Hartley et al., 2017) at https://climate.esa.int/en/data; WWF terrestrial ecoregions (Olson et al., 2001) at https://www.worldwildlife.org/publications/terrestrial-ecoregions-of-the-world; and the trait products of Dong et al. (2023), van Bodegom et al. (2014), Butler et al. (2017), Madani et al. (2018), Moreno-Martínez et al. (2018), and Boonman et al. (2020), which were obtained from the supplement of Dong et al. (2023).

## Author contributions

Conceptualization: LL, TH. Methodology: LL. Software: LL, SS (see also Acknowledgements). Formal analysis: LL with comments from all authors. Investigation: LL. Resources: Senckenberg Biodiversity and Climate Research Centre provided computational resources. Data curation: LL. Writing – original draft preparation: LL and TH drafted the manuscript with contributions from all authors. Writing – review and editing: all authors. Visualization: LL, TA, MDP, TH with contributions from all authors.

## Appendix A

**Figure A1.**
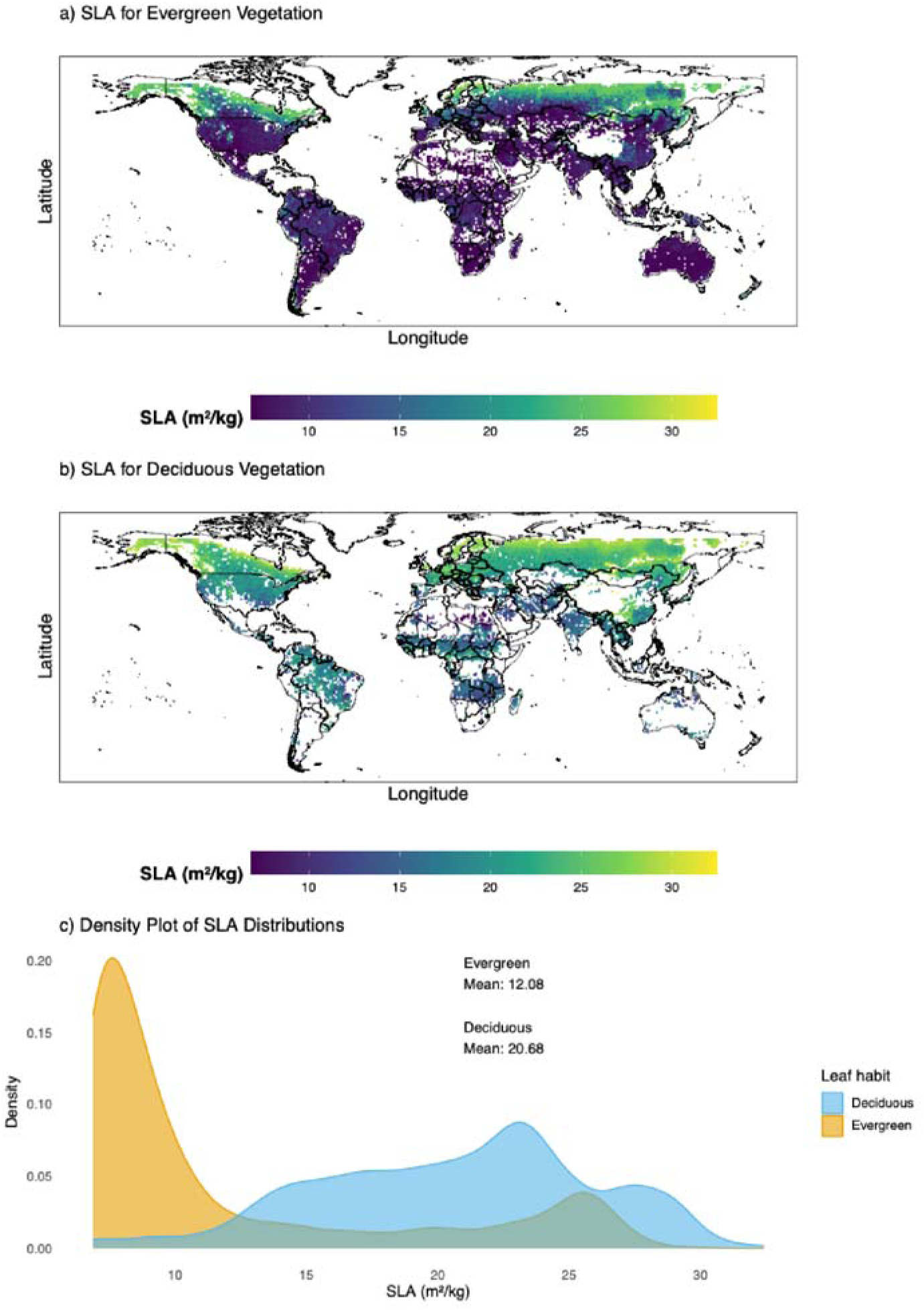
Global distribution of aDGVM2 simulated SLA. (a) Spatial distribution of SLA for evergreen leaf phenology. (b) Spatial distribution of SLA for deciduous leaf phenology. (c) Density plots of SLA (m^2^ kg^−1^).

**Figure A2.**
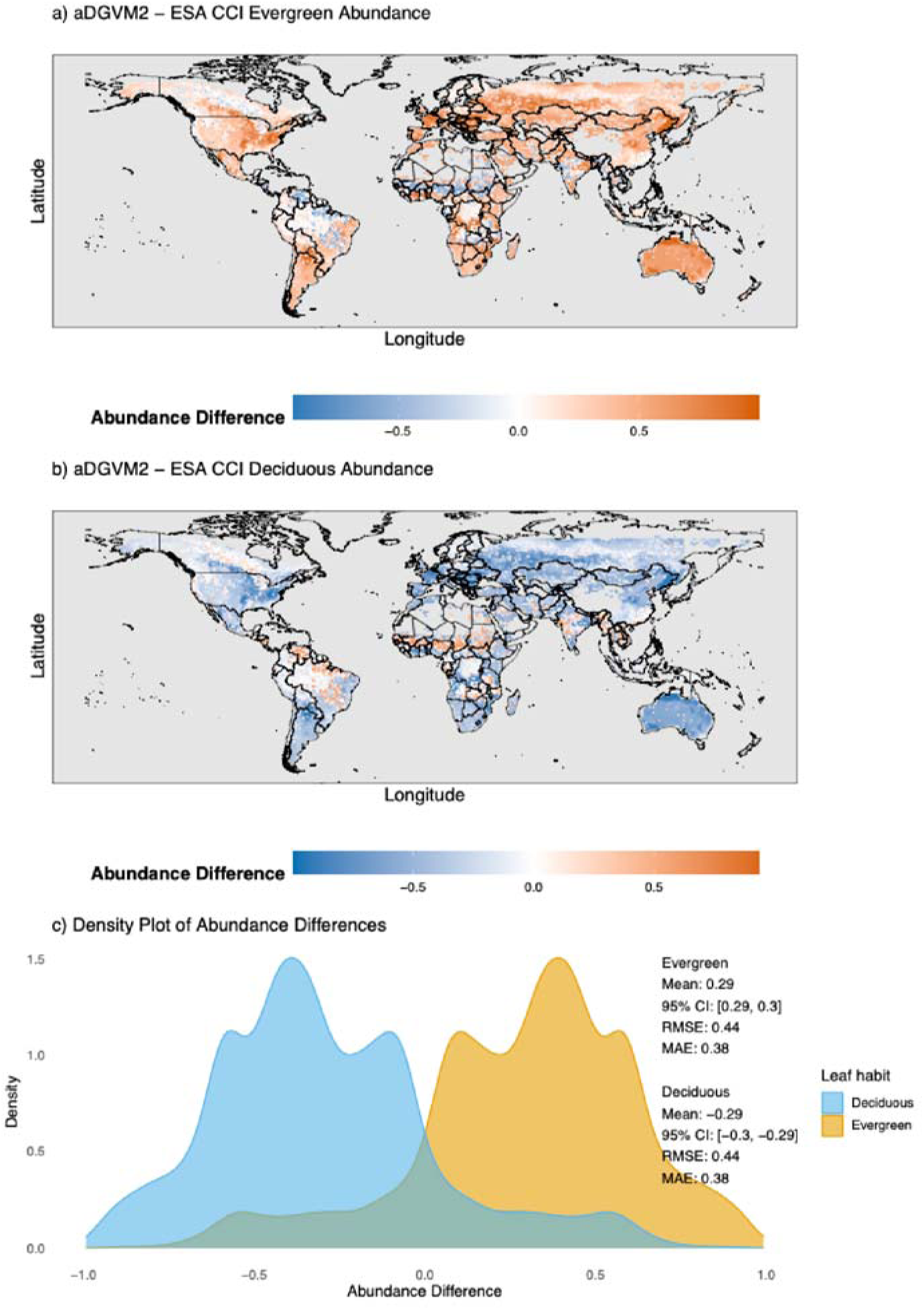
Global differences in leaf phenology between aDGVM2 and ESA-CCI-derived abundances. (a) Spatial distribution of evergreen differences. (b) Spatial distribution of deciduous differences. (c) Density plots of abundance differences.

## Acknowledgements

Steven Higgins contributed to developing the model and plant hydraulics sub-modules. LL wrote the draft manuscript and used grammar correction software, which included a generative AI interface, to refine the text.

